# Major components in the KARRIKIN INSENSITIVE2-ligand signaling pathway are conserved in the liverwort, *Marchantia polymorpha*

**DOI:** 10.1101/2020.11.17.387852

**Authors:** Yohei Mizuno, Aino Komatsu, Shota Shimazaki, Xiaonan Xie, Kimitsune Ishizaki, Satoshi Naramoto, Junko Kyozuka

## Abstract

KARRIKIN INSENSITIVE2 (KAI2) was first identified in *Arabidopsis thaliana* as a receptor of karrikin, a smoke-derived germination stimulant. KAI2 is also considered a receptor of an unidentified endogenous molecule called the KAI2-ligand (KL). Upon KAI2 activation, signals are transmitted through degradation of D53/SMXL proteins via ubiquitination by a Skp-Cullin-F-box (SCF) E3 ubiquitin ligase complex. All components in the KL signaling pathway exist in the liverwort *Marchantia polymorpha*, namely Mp*KAI2A* and Mp*KAI2B*, Mp*MAX2* encoding the F-box protein, and Mp*SMXL*, indicating that the signaling pathway became functional in the common ancestor of bryophytes and seed plants. Genetic analysis using knock-out mutants of these KL signaling genes, produced using the CRISPR system, indicated that Mp*KAI2A*, Mp*MAX2* and Mp*SMXL* act in the same genetic pathway and control early gemma growth. Introduction of Mp*SMXL^d53^*, in which a domain required for degradation is mutated, into wild-type plants caused phenotypes resembling those of the Mp*kai2a* and Mp*max2* mutants. In addition, Citrine fluorescence was detected in tobacco cells transiently transformed with the *35S*:Mp*SMXL*-*Citrine* gene construct and treated with MG132, a proteasome inhibitor. On the other hand, introduction of *35S*:Mp*SMXL^d53^*-*Citrine* conferred Citrine fluorescence without MG132 treatment. These findings imply that MpSMXL is subjected to degradation, and that degradation of MpSMXL is crucial for KL signaling in *M. polymorpha*. We also showed that MpSMXL is negatively regulated by KL signaling. Taken together, this study demonstrates that basic mechanisms in the KL signaling pathway are conserved in *M. polymorpha*.

## Introduction

Plants alter their developmental program to optimize their growth depending on environmental conditions. Plant hormones are crucial, linking environmental conditions with plant growth. Strigolactones (SLs), carotenoid-derived terpenoid lactones, are unique molecules that act as plant hormones *in planta* and control variable aspects of plant growth, including shoot branching, root growth and senescence (1, 2). In addition, they act as rhizosphere-signaling molecules in the soil when exuded from roots (3). Here, they promote symbiosis with arbuscular mycorrhiza (AM), facilitating phosphate acquisition from soil. In seed plants, SL is perceived by the α/β-hydrolase superfamily protein, DWARF14 (D14) (4–8). D14 is part of a small gene family containing a few to several genes that are divided into two major groups, one containing *D14* and the other containing *KARRIKIN INSENSITIVE2* (*KAI2*) (9–11).

KAI2 was first identified in the model plant species, Arabidopsis, as a receptor of karrikin, a butenolide molecule found in smoke from burned vegetation (9, 11). Karrikins play a role in the regulation of seed germination and seedling development. However, because loss-of-function mutants of *KAI2* showed smoke-independent phenotypes, KAI2 was proposed to be a receptor of an unidentified endogenous molecule called KAI2 ligand (KL) (9, 12–14). Although the detailed mechanisms involved remain to be elucidated, it is widely accepted that the SL and KARRIKIN/KL signals are transduced by parallel proteolysis-dependent signaling pathways in which D14 and KAI2 work as their receptors, an F-box protein, which forms a Skp-Cullin-F-box (SCF) E3 ubiquitin ligase complex with Skp1 (ASK1) and Cullin (CUL1), and repressor proteins, are major components (15–17). DWARF3 (D3) of *Oryza sativa* (rice) and MAX2 of Arabidopsis are F-box proteins and are shared in both the SL and KL signaling pathways (18–20). The D53 protein of rice was identified as a degradation target of SL signaling (15, 17). SMAX1 and SMAX1-LIKE2 (SMXL2) of Arabidopsis, close homologs of D53, were identified as likely targets of KL signaling (16, 21, 22). In rice, perception of SLs promotes a physical interaction of D14 with D3 and D53, which leads to rapid degradation of D53, resulting in activation of downstream growth responses (15, 17). The *D53* and *SMAX1* genes form a small gene family (23–25). Among the eight *SMXL* genes of Arabidopsis, *SMXL6*, *7* and *8* are redundant orthologs of *D53*, and are involved in the control of shoot branching regulated by SL signaling (16). *SMAX1* and *SMXL2* are involved in the KAI2-dependent pathway controlling seed germination and hypocotyl elongation (14, 21, 26). Recently, it was shown that proteolysis of SMXL2 can be triggered by both SLs and Karrikins (KARs) (27). On the other hand, SMXL3, 4 and 5 have lost the sequence required for degradation during evolution and are thus not under proteolysis-dependent regulation (28). It has been proposed that SMXL6, 7, 8 repress gene expression through interaction with transcription factors and/or transcriptional corepressors. In addition, SMXL6 of Arabidopsis directly binds DNA and regulates transcription of downstream genes (27). These findings suggest that the expansion of the D53/SMXL protein families was likely to be important in conferring diversity in the function of SL and KL signaling, enabling control of many aspects of growth and development.

Land plants evolved from an ancestral charophycean alga more than 450 million years ago (29, 30). Basal land plants and green algae contain at least one *KAI2* ortholog while *D14* orthologs exist only in seed plants (9, 10, 31). This indicates that *KAI2 is* ancestral and *D14* arose by gene duplication events before the evolution of seed plants. Among the components in the SL and KL signaling pathways, the D3/MAX2 F-box proteins are highly conserved in all embryophytes and Charales including Coleochaetales (10), whereas the D53/SMXL family proteins are present in all major land plant groups, but are not identified in charophytes (23–25). Therefore, an entire set of major KL signaling components are aligned in bryophytes although their function is poorly understood. *Marchantia polymorpha*, the common liverwort, is suitable for molecular genetic studies because of its low genome redundancy, and the availability of its whole genome sequence and essential tools for molecular genetic studies (29). Moreover, the existence of a single copy of the *D53/SMXL* gene in *M. polymorpha* makes it a highly suitable species to understand the ancestral function and prototype of this signaling pathway, as well as allowing us to trace the evolution of this pathway. Here, as a first step toward understanding the ancestral role and evolution of SL and KL signaling in plants, we analyzed the function of genes putatively involved in this signaling in *M. polymorpha*.

## Results

### Genes in the KL signaling pathways in *Marchantia polymorpha*

In this paper, we refer to the putative signaling pathway that supposedly mediates signaling derived from the KAI2-ligand (KL) as the KL signaling pathway. Genetic nomenclature is as outlined in (32). Two *KAI2* orthologs, one *D3/MAX2* ortholog and one *D53/SMXL* ortholog (termed Mp*KAI2A* and Mp*KAI2B*, Mp*MAX2* and Mp*SMXL*, respectively) have been identified in the genome of *M. polymorpha* (29, 33). KAI2 and D14 contain three conserved amino acids that are crucial to their hydrolase activity (8, 10, 34). These three amino acids form a catalytic triad that is conserved in MpKAI2A and MpKAI2B (*SI Appendix*, Fig S1 *A*). MpMAX2 contains a conserved C-terminal α-helix that is required in D3 for interaction with D14 and to recruit D53 (*SI Appendix*, Fig S1 *B*; 35). Amino acids conserved in SMXL proteins, including the RGKT motif required for degradation and the ethylene response factor-associated amphiphilic repression (EAR) motif required for repression of transcription in other species, are also conserved in MpSMXL (*SI Appendix*, Fig S1 *C*; 24). We first examined the tissue specificity of the expression of Mp*KAI2A*, Mp*KAI2B*, Mp*MAX2* and Mp*SMXL*, and confirmed that all four genes were widely expressed in the thallus, gemma, antheridiophore and archegoniophore, and the male and female gametophores, respectively (*SI Appendix*, Fig S2). Then, loss-of-function mutants of these genes were generated in *M. polymorpha* using a CRISPR/Cas9 genome editing system (36). Mutations in the transgenic plants were confirmed by sequencing and at least two independent alleles for each gene, representing complete loss-of-function alleles, were selected for subsequent phenotypic analysis (*SI Appendix*, Fig S3).

### Mp*KAI2A* and Mp*MAX2* work in the same genetic pathway in the control of thallus growth

We observed thallus growth in mutants of three putative KL signaling genes (Fig. 1, *SI Appendix*, Fig. S4). At least two independent alleles of each gene were analyzed and the results of the second alleles are shown in *SI Appendix*, Fig. S4. Thalli from mutant plants with either allele of Mp*kai2b* (Mp*kai2b-1* or Mp*kai2b-2*) were morphologically indistinguishable from those of wild-type (WT) plants (Fig. 1 *A-F*). In contrast, thalli from mutant plants with either allele of Mp*kai2a* (Mp*kai2a-1* or Mp*kai2a-2*) or Mp*max2* (Mp*max2-1* or Mp*max2-2*) showed similar alterations, in which early growth of the thalli was retarded and the thalli curved upward. The size of the thalli of Mp*kai2a* and Mp*max2* was significantly smaller than that of the WT whereas there was no significant difference between Mp*kai2a* and Mp*max2* (Fig. 1 *C* and *D*). The angle of thallus curving was larger in Mp*kai2a* and Mp*max2* plants compared to the WT, while no significant differences were observed between Mp*kai2a* and Mp*max2* (Fig. 1 *E* and *F*). To test the possibility that the defects in the Mp*kai2b* mutants were masked by the function of the Mp*KAI2A* gene, which contains a high sequence similarity to Mp*KAI2B*, we produced double mutants of Mp*KAI2a* and Mp*KAI2b*. The Mp*kai2a* Mp*kai2b* double mutants were not significantly different from Mp*kai2a*, implying that Mp*KAI2b* is not significantly involved in the control of thallus growth (Fig. 1 *C-F*). These data indicate that Mp*KAI2A* and Mp*MAX2* control thallus growth in a similar way.

**Fig. 1.**
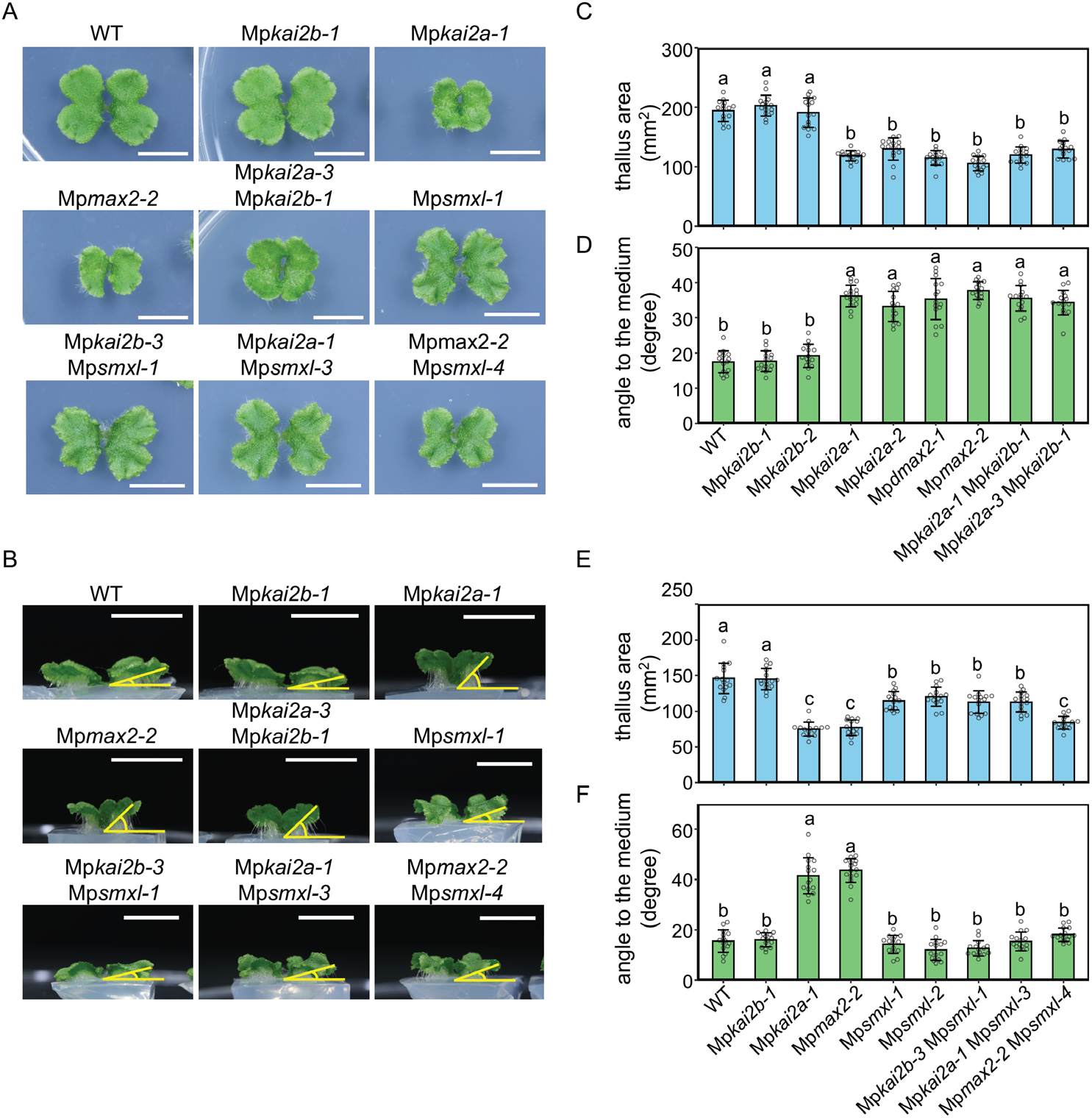
Growth phenotypes of loss-of-function mutants of KL signaling genes. **(A)** Top view of 14-day-old thalli. Bars = 1 cm. **(B)** Side view of the thalli shown in (A). Bars = 1 cm. **(C, E)** Area of 14-day-old thalli. **(D, F)** Angles between the 14-day-old thalli and the growth media. Data in (C) to (F) are the mean ± SD from three independent analyses. Fifteen gemmae were used in each analysis. Tukey’s honestly significant difference (HSD) test was used for multiple comparisons. *P*-values < 0.05 were considered statistically significant.

### Mutations in Mp*SMXL* suppress defects in Mp*kai2a* and Mp*max2* mutants

*M. polymorpha* contains a single SMXL/D53 family gene, Mp*SMXL*. The domains that are conserved in the SMXL/D53 family proteins in Angiosperms, including the RGKT motif required for its SCF^D3/MAX2^-dependent degradation and the EAR motif required for transcriptional repression of downstream genes, are also conserved in MpSMXL (*SI Appendix*, Fig. S1 *C*; 24, 29). In Mp*smxl* loss-of-function mutants, the size of the thallus was reduced to a size between that of the WT and the Mp*kai2a* or Mp*max2* plants (Fig. 1 *A* to F; *SI Appendix*, Fig. S4). In contrast to Mp*kai2a* and Mp*max2*, Mp*smxl* mutants did not show upward curving of the thalli (Fig. 1 *B* and *F*). The morphology and the size of the thalli in the Mp*smxl* Mp*kai2a* and Mp*smxl* Mp*kai2b* double mutant plants was not significantly different from that of the Mp*smxl* single mutant plants. The thallus-curving phenotype of Mp*kai2a* and Mp*max2* plants was suppressed in the double mutants of Mp*smxl* Mp*kai2a* and Mp*smxl* Mp*max2*, indicating that Mp*SMXL* works downstream of Mp*KAI2a* and Mp*MAX2* (Fig. 1 *C* and *D*). However, the size of the thalli in Mp*smxl* Mp*max2* double mutant plants was slightly smaller than that of the Mp*smxl* plants (Fig. 1 *A* and *D*). This may indicate that SMXL works somewhat independently from Mp*MAX2*.

To investigate the cause of the reduced thallus size, we examined the gemmae in more detail, because many gemmae of a uniform developmental stage can be easily obtained and it is possible to count the number of cells due to their small size and flat shape (Fig. 2 *A*). We measured the size of the gemmae and counted the total number of their epidermal cells (Fig. 2 *B* and *C*). Gemmae in Mp*kai2a*, Mp*max2* and the Mp*smxl* mutant plants were smaller than those of the WT plants at the mature gemma stage. Consistent with this, the number of cells in the gemmae was reduced in Mp*kai2a* and Mp*max2* plants (Fig. 2 *C*). On the other hand, despite the decrease in gemmae size, the number of cells in the gemmae was increased in Mp*smxl* plants (Fig. 2 *C*). Mp*kai2a* Mp*smxl*, Mp*kai2b* Mp*smxl* and Mp*max2* Mp*smxl* double mutants also contained more cells in the gemma than in the WT plants. Cell size estimated from the gemma area and the number of cells was not significantly different between WT, Mp*kai2a*, Mp*kai2b* and Mp*max2* mutants (Fig. 2 *D*). In contrast, the estimated cell size was smaller in Mp*smxl* and double mutants containing Mp*smxl* compared to that of the WT plants. These results indicate that Mp*KAI2A* and Mp*MAX2* act to increase the cell number in the gemmae without affecting the cell size, while Mp*SMXL* functions to suppress cell proliferation and to promote cell growth. In addition, these results support the hypothesis that Mp*SMXL* works downstream of Mp*KAI2A* and Mp*MAX2* in the genetic pathway which regulates gemma development.

**Fig. 2.**
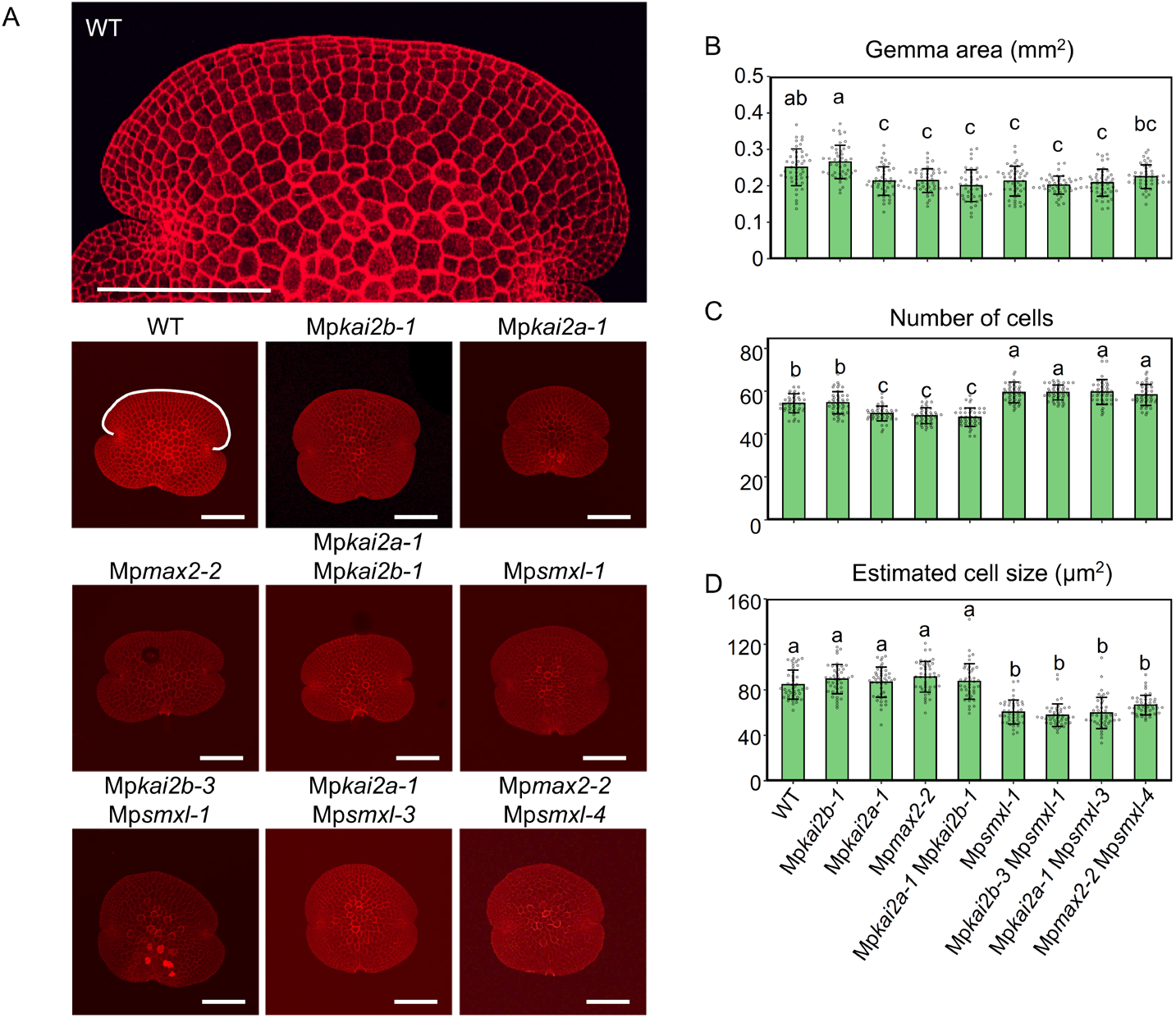
Phenotypes in gemma size and cell number. **(A)** Gemmae of WT and mutant plants stained with propidium iodide. Bars = 200 μm **(B)** Area of gemmae. **(C)** Number of cells in the outermost cell layer of the dorsal side of gemmae as shown outlined in white on the image of the WT plant in (A). **(D)** Relative size of a cell estimated from the gemma area shown in (B) and the number of cells shown in (C). Data shown in (B) and (C) are the mean ± SD from three independent experiments. Fifteen gemmae were used in each experiment. The HSD test was used for multiple comparisons. *P*-values < 0.05 were considered statistically significant.

### Degradation of MpSMXL is required in the signaling pathway through MpKAI2A and MpMAX2

We next analyzed whether degradation of MpSMXL is required in the KL signaling pathway in *M. polymorpha* as is the case for the SMXL/D53 family proteins in rice and Arapidopsis (15–17, 21, 22). In rice and Arabidopsis, D53/SMXL proteins containing the RGKT motif are ubiquitinated via the SCF^D3/MAX2^ complex and degraded through the 26S proteasome.

To observe the MpSMXL protein, MpSMXL was fused with Citrine, and driven by its own promoter in *M. polymorpha*. We produced more than 10 independent transgenic lines and confirmed expression of the introduced MpSMXL-Citrine mRNA, however, we did not detect Citrine fluorescence under the microscope, nor did we detect MpSMXL-Citrine protein by Western blotting in any of the transgenic lines, even after treatment with MG132, a proteasome inhibitor. This suggested the possibility of proteasome-dependent degradation of the MpSMXL-Citrine protein. To test this, we used a transient expression system in tobacco (*Nicotiana benthamiana*) epidermis cells (Fig. 3). The Mp*SMXL* gene fused with the Citrine gene and the CaMV35S promoter was transformed into tobacco cells. Citrine fluorescence was detected in the tobacco cells after treatment with MG132, supporting our hypothesis that MpSMXL is degraded (Fig. 3 *A* and *B*). To further confirm this, we replaced the Mp*SMXL* gene with the degradation-resistant Mp*SMXL^d53^*, in which the RGKT domain was mutated to the same sequence as the degradation-resistant *d53* mutant allele of rice (15–17, 26), fused with the Citrine gene and transformed into tobacco cells. Citrine fluorescence was observed without MG132 treatment (Fig. 3 *C*). These results indicate that MpSMXL is subjected to proteosome-dependent degradation.

**Fig. 3.**
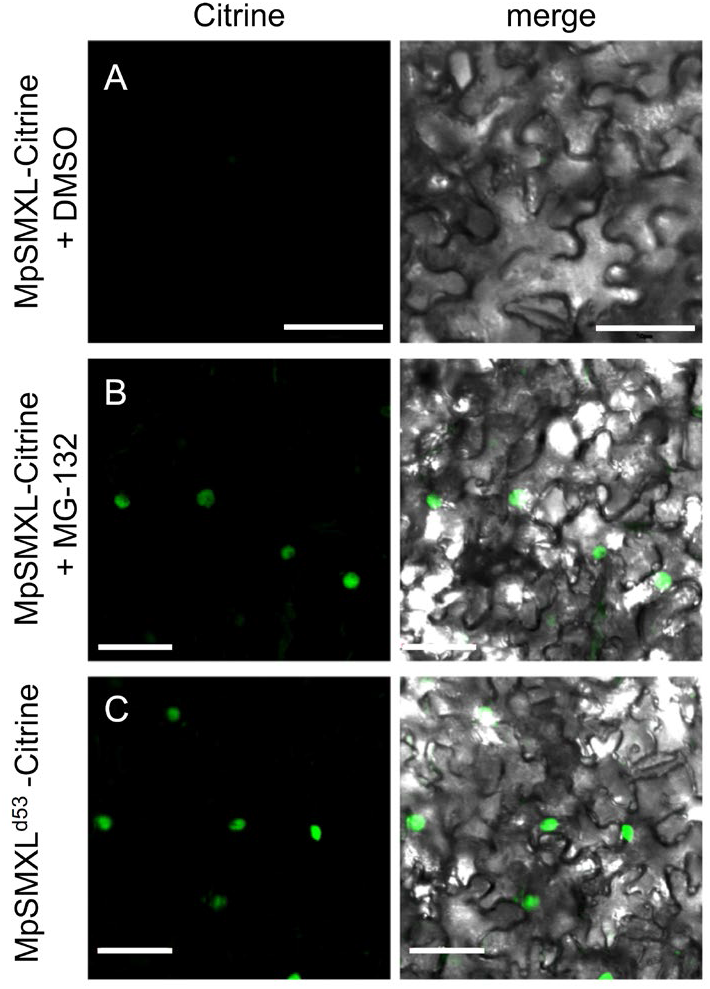
Localization of MpSMXL-Citrine in *N. benthamiana* epidermal cells. Subcellular localization of MpSMXL-Citrine in *N. benthamiana* epidermal cells treated with 0.25% (v/v) DMSO **(A)**, or 100 μM MG-132 **(B)**. Subcellular localization of MpSMXL^d53^-Citrine in *N. benthamiana* epidermal cells **(C)**. Micrographs showing cells expressing MpSMXL-Citrine or MpSMXL^d53^-Citrine were examined under fluorescence (left) and bright field (right) by confocal microscopy. Bars = 50 μm.

To understand the consequence(s) of degradation on the function of MpSMXL, the Mp*SMXL^d53^* gene was fused with the promoter region of Mp*SMXL* and transformed into WT *M. polymorpha*. We generated more than twenty independent transgenic lines (*_pro_SMXL:SMXL^d53^*). Although the severity of the phenotype varied among the transgenic lines, most showed defects resembling Mp*kai2a* and Mp*max2*, in which the thallus is small and curved upward. We selected transgenic lines showing moderate (*_pro_SMXL:SMXL^d53^* −10, −12) and strong (*_pro_SMXL:SMXL^d53^* −3, −9) phenotypes for further analysis (Fig. 4). The lines with strong phenotypes showed equivalent defects to those of the Mp*kai2a* and Mp*max2* mutant plants (Fig. 4 *A-C*). These data suggest that the degradation of MpSMXL is required in KL signaling.

**Fig. 4.**
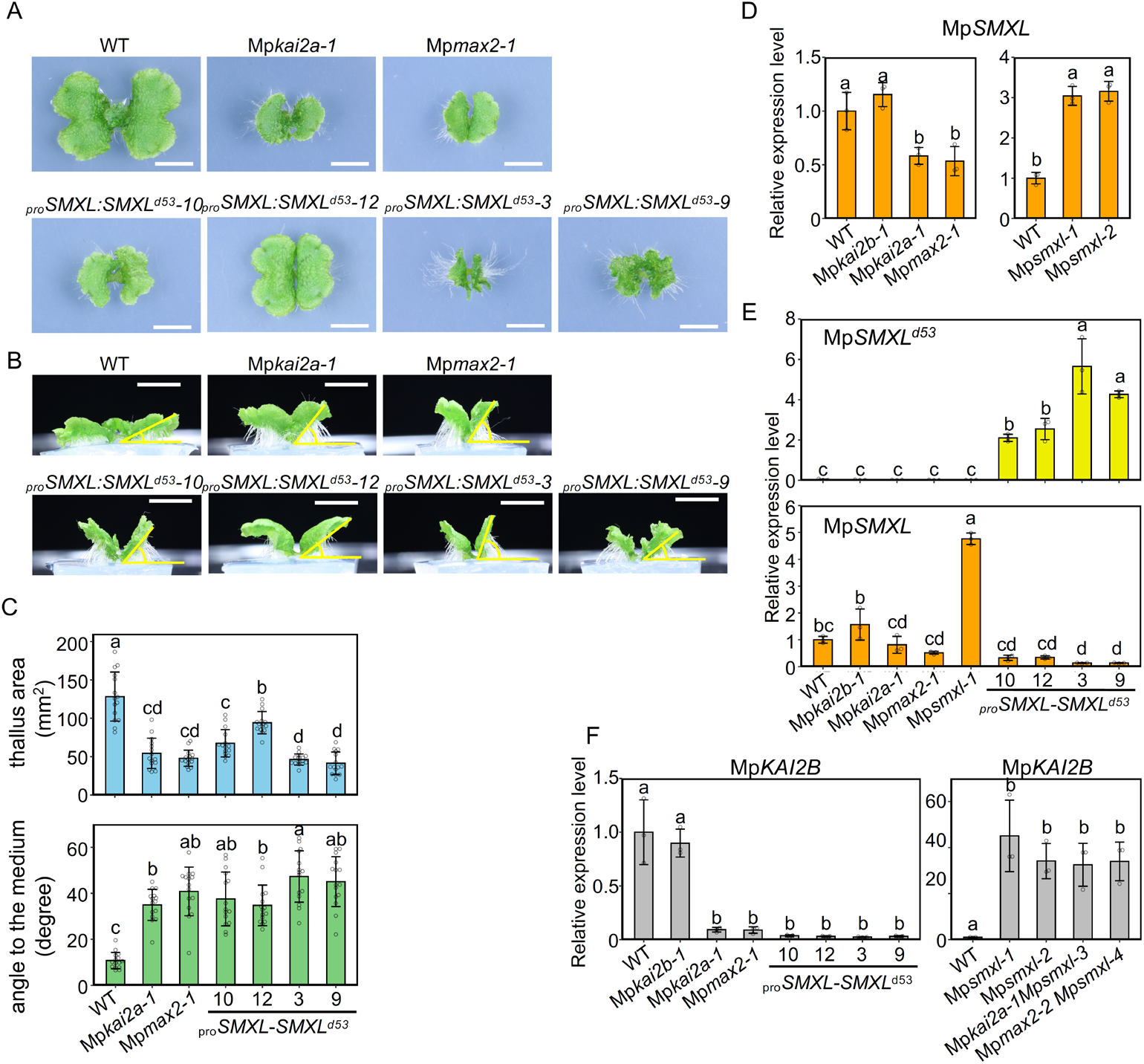
Functional analysis of the conserved RGKT motif of MpSMXL. **(A)** to **(C)** Effects of expression of the Mp*SMXL* gene containing the same mutation as rice *d53* (*_pro_SMXL:SMXL^d53^*) on thallus growth. Top (A) and side (B) view of 14-day-old thalli. Bars = 1 cm. **(C)** Thallus area and angle to the growth medium. Data are the mean ± SD from three independent analyses. Fifteen gemmae were used in each analysis. **(D)** Expression of the Mp*SMXL* gene relative to Mp*ELONGATION FACTOR* (Mp*EF*) in the thallus. **(E)** Expression of the introduced *_pro_SMXL:SMXL^d53^* and endogenous Mp*SMXL* genes relative to Mp*EF* in the thallus. **(F)** Expression of Mp*KAI2B* relative to Mp*EF* in the thallus. Data in (D) and (F) are the mean ± SD from three biological replications. The HSD test was used for multiple comparisons in (C), (D) and (F). *P*-values < 0.05 were considered statistically significant.

We next examined feedback regulation in the expression of KL signaling genes. In rice and Arabidopsis, accumulation of *D53* and *SMXL678* transcripts is subjected to negative feedback control in the SL signaling pathway (15, 17). In rice, *D53* expression is suppressed in SL signaling and biosynthesis mutants while it is upregulated by the addition of GR24, a synthetic SL. In *M. polymorpha*, the expression level of Mp*SMXL* mRNA was decreased in the Mp*kai2a* and Mp*max2* mutants while it was enhanced in Mp*smxl* loss-of-function mutants (Fig. 4 *D*). This implies that Mp*SMXL* expression was subjected to conserved feedback suppression by KL signaling.

In the transgenic lines, the level of introduced *_pro_SMXL:SMXL^d53^* expression was roughly consistent with the severity of the defects (Fig. 4 *E*). The amount of endogenous Mp*SMXL* transcript, however, appeared to decrease depending on the severity of the defects (Fig. 4 *E*). Moreover, we found that Mp*KAI2B* expression was dramatically reduced in Mp*kai2a* and Mp*max2* plants (Fig. 4 *F*). This implies that Mp*KAI2B* expression is dependent on KL signaling. On the other hand, Mp*KAI2B* expression was dramatically enhanced in Mp*smxl-1*, the loss-of-function mutant of Mp*SMXL*. This further supports the hypothesis that Mp*KAI2A*, Mp*MAX2* and Mp*SMXL* work in the same pathway. We observed that Mp*KAI2B* expression was severely suppressed in moderate (−10 and −12) and strong (−3 and −9) *_pro_SMXL-SMXL^d53^* lines, indicating that KL signaling was dampened by the suppression of MpSMXL protein degradation in the transgenic lines. All these results clearly indicate that MpSMXL works as the repressor in the MpSMXL degradation-dependent KL signaling pathway, in which MpKAI2A and MpMAX2 are involved.

### *M. polymorpha* does not respond to karrikins

To obtain insights into the ligands involved in the KL signaling pathway in *M. polymorpha*, we examined the responses to karrikins (KAR1 and KAR2) and the synthetic strigolactone, *rac*-GR24. Gemmae were grown on media containing *rac*-GR24, KAR1 or KAR2 and their growth was measured after fourteen days’ culture. Growth of thalli was not affected by addition of 2 μM *rac*-GR24 while it was severely suppressed by the addition of a high concentration (5 μM) of *rac*-GR24 (Fig. 5 *A*, *SI Appendix*, Fig. S5 *A*). The suppression was weakened in Mp*kai2a-1* and Mp*max2-2* and in the Mp*kai2a-1* Mp*max2-1* double mutants, while it was not affected in the Mp*kai2b-1* plants. On the other hand, addition of neither KAR1 nor KAR2 affected the growth of thalli in any of the genotypes examined. We asked whether expression of Mp*KAI2B* and Mp*SMXL*, markers that are positively and negatively regulated by the KL signaling pathway, respectively (Fig. 5*D*), is changed by addition of the chemicals. Neither Mp*KAI2B* nor Mp*SMXL* expression was affected by the addition of a high concentration (10 μM) of *rac*-GR24. This suggests that the growth retardation caused by the addition of a high concentration (higher than 5 μM) of *rac*-GR24 is partially dependent on Mp*MAX2* but independent of KL signaling. Finally, we investigated the physical interaction between MpKAI2A or MpKAI2B with four stereoisomers of GR24 by Differential Scanning Fluorimetry analysis. Only (-)-GR24, that has an unnatural D-ring configuration, interacted slightly with MpKAI2A and MpKAI2B (*SI Appendix*, Fig. S5 *B*).

**Fig. 5.**
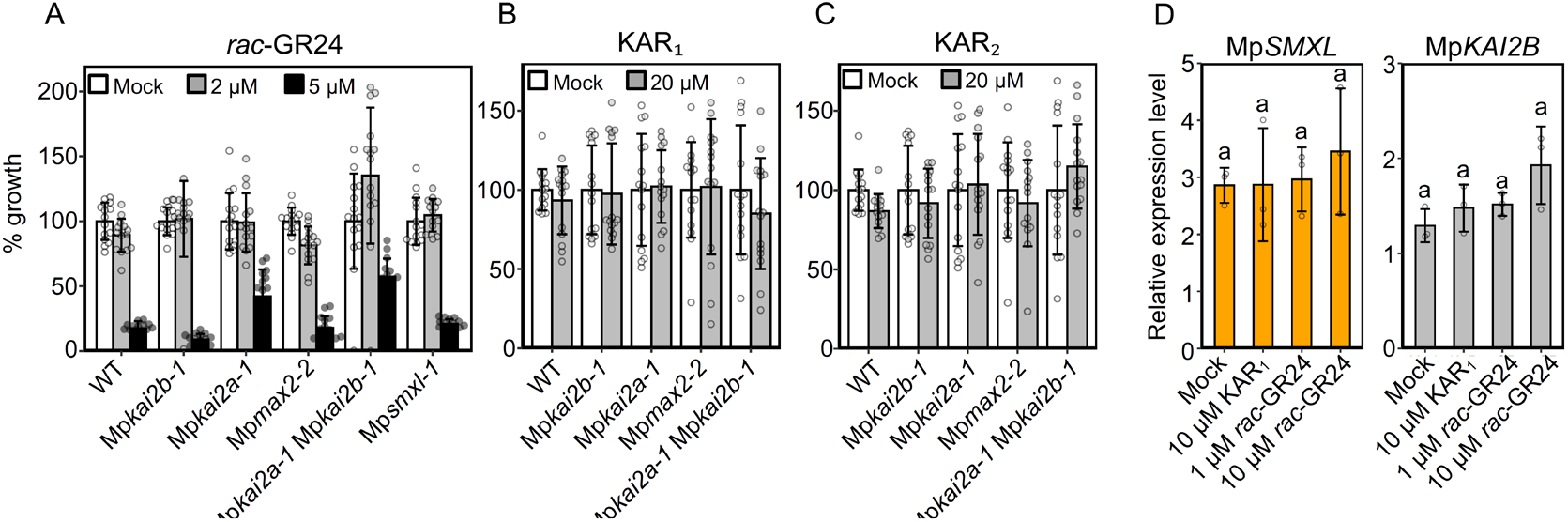
Effects of exogenously supplied *rac*-GR24, KAR1 and KAR2 on thallus growth. **(A)** to **(C)** Growth of thalli exposed to 2 or 5 μM *rac*-GR24 **(A)**, 20 μM KAR1 **(B)** and 20 μM KAR2 **(C)**, relative to a mock-treated control. Fifteen gemmae were used in each experiment. Data are the mean ± SD from three independent experiments. **(D)** Expression of Mp*SMXL* and Mp*KAI2B* relative to Mp*EF* in the thalli exposed to 10 μM KAR1 and 1 μM or 10 μM *rac*-GR24.

## Discussion

### Components in the KL signaling pathway are conserved in *M. polymorpha*

The *M. polymorpha* genome contains two *KAI2*, one *D3/MAX2* and one *SMXL* homologs (29). In this study, we showed that loss-of-function mutants of Mp*KAI2A*, one of the two *KAI2* homologs, and Mp*MAX2*, showed indistinguishable defects in gemma and thallus growth and that Mp*smxl* loss-of-function mutants were epistatic to Mp*kai2a* and Mp*max2*. These results indicate that Mp*KAI2A*, Mp*MAX2* and Mp*SMXL* function in the same genetic pathway.

In the dominant *d53* mutant of rice, an arginine in the RGKT motif is replaced with threonine and a subsequent GKTGI amino acid sequence is deleted. This mutation confers resistance to degradation of the D53 protein upon SL perception and leads to more branched phenotypes in rice (15–17). Deletion of the RGKT sequence from Arabidopsis SMXL6 also confers resistance to degradation and causes more shoot branching phenotypes (16). We showed that introduction of the mutant version of the Mp*SMXL* gene containing the same amino acid sequence as the *d53* mutant of rice into WT *M. polymorpha* mimicked the phenotypes of the Mp*kai2a* and Mp*max2* mutants. We also showed that introduction of the Citrine gene fused with MpSMXL into tobacco epidermis cells resulted in Citrine fluorescence when the introduced MpSMXL contained the *d53* mutation or the tobacco cells were treated with proteolysis inhibitor. All these findings support our hypothesis that degradation of MpSMXL is crucial in the signaling of MpKAI2A.

F-box protein-mediated protein degradation is used for signaling of multiple plant hormones and was established in basal land plants (9, 37). In *M. polymorpha* and *Physcomitrella patens*, another bryophyte, auxin is perceived by TIR1, an F-box protein, and the signal is transduced through degradation of AUX/IAA transcription factors, as is the case in seed plants (38–41). The F-box protein coronatine-insensitive 1 (COI1), the receptor of jasmonate in seed plants, works as a receptor of dinor-12-oxo-phytodienoic acid (dinor-OPDA), a molecule related to jasmonate, in the proteolysis-dependent signaling cascade in *M. polymorpha* (42). Although the F-box protein ancestral to TIR and COI1 exists in charophytes, the sequences necessary for interaction with ligands and repressor proteins are lacking, suggesting that TIR1 and COI1 established their function by neofunctionalization during the evolution of basal land plants (43). *D3/MAX2* orthologs but not *SMXL* are present in charophytes. Taking all these observations into account, it appears that the proteolysis-dependent signaling pathway through KAI2 and D3/MAX2 was established in the common ancestor of bryophytes and seed plants.

The number of *SMXL* genes has increased during evolution. The multiple copies of *SMXL* genes in seed plants are classified into subgroups, each regulating different aspects of growth and development (21, 22, 26, 28, 44). In contrast, *M. polymorpha* contains only a single *SMXL* gene, which functions in the MpKAI2A signaling pathway. We cannot exclude the possibility that the MpKAI2A and MpKAI2B pathways share MpSMXL as a repressor, however, the fact that Mp*SMXL* expression is not affected in Mp*kai2b* indicates that MpSMXL is unlikely to be involved in the signaling of MpKAI2B. It is of interest to know whether the function of MpSMXL fully depends on MpKAI2A and MpMAX2 or whether MpSMXL acts partially independently from them.

### Possible ligands of MpKAI2A and MpKAI2B

The D14 clade, including core D14 genes and D14 Like 2 (DLK2), arose through gene duplication of KAI2s (10). In Arabidopsis, D14 specifically recognizes SLs and KAI2 does not respond to natural SLs (9, 12). Based on this, it is hypothesized that generation of D14 led to the establishment of the SL perception system (45). The existence of canonical SLs in basal land plants is currently under debate (31, 46–49). In *P. patens*, protonema development is affected in carotenoid cleavage dioxygenase 8 (*ccd8*) mutants, in which SL biosynthesis is blocked, and the defects are complemented by application of GR24, indicating that the products of CCD8 or their derivatives are active (48). However, MAX1 is absent in *P. patens*, therefore the active molecule responsible for this trait may be different from canonical SLs. If basal land plants produce SLs, how they are perceived without D14 remains unknown. Recently, it was found that the KAI2 genes are extensively amplified in parasitic plants and some of them act as highly sensitive SL receptors (50, 51). This implies that the machinery for SL perception can be generated through modification of ligand reception specificity of KAI2 so that they can respond to SLs. These findings raise the possibility that a subset of KAI2s may work as SL receptors in basal land plants, while their function as SL receptors was substituted by D14 in seed plants (52).

It is unlikely that SLs are produced in *M. polymorpha*, at least through a canonical biosynthesis pathway, due to the absence of CCD8 and MAX1 (29). This suggests that the ligand of MpKAI2A is probably not an SL. Indeed, MpKAI2A driven by the Arabidopsis KAI2 promoter did not complement the shoot branching phenotype of Arabidopsis *d14* mutants, indicating that MpKAI2A does not respond to Arabidopsis endogenous SLs (33). We showed that thalli responded to *rac*-GR24 in a manner partially dependent on MpKAI2A and MpMAX2. It is known that AtKAI2 responds to GR24^ent-5DS^, a synthetic analog of a non-natural compound (33, 53). Results of Differential Scanning Fluorimetry (DSF) analysis in this study indicated that MpKAI2A and MpKAI2B probably interact with GR24^ent-5DS^. The closest homologs of MpKAI2A in *P. patens*, PpKAI2L-C, -D and -E, show a shift in the protein melting point with high specificity to ent-5-deoxystrigol, a non-natural strigolactone isomer (54). Considering these data, the ligand specificity of MpKAI2A may be similar to that of AtKAI2 and the natural ligand of MpKAI2A may be KL or a KL-related molecule. However, MpKAI2A expressed by the Arabidopsis KAI2 promoter did not rescue the seedling phenotype of *kai2-2*, suggesting the possibility that KLs have diverged between Arabidopsis and *M. polymorpha* (33). The absence of a response to exogenously supplied KAR1 or KAR2 observed in this study also supports the idea that the specificity of ligand perception has also diverged in seed plants. However, in cross-species complementation experiments (33), the possibility that MpKAI2A does not physically interact with Arabidopsis SMAX1, SMXLs or MAX2 cannot be ruled out.

### Function of MpKAI2B

We did not find any defects in growth in loss-of-function Mp*kai2b* mutants even though Mp*KAI2B* is expressed in all organs examined at a comparable level to that of Mp*KAI2A*. A possible explanation for the lack of a Mp*kai2b* phenotype is that MpKAI2B works as an SL receptor in *M. polymorpha*, however, due to the loss of SL biosynthesis ability in *M. polymorpha*, the function of MpKAI2B is masked. However, the fact that we did not detect a physical interaction between GR24^5DS^ or GR24^4DO^, synthetic analogs of natural compounds, and MpKAI2A or MpKAI2B does not support this possibility. Alternatively, MpKAI2B may function under special conditions or in specific cells and defects are not apparent under normal growth conditions. To reveal the function of MpKAI2B, a more detailed and comprehensive analysis of loss-of-function mutants under variable growth conditions using other experimental tools, such as transcriptomics, proteomics and metabolomics, is required. Although the lack of clear phenotypes in the loss-of-function mutants has hampered our understanding of its function, *DLK2* is used as a positive marker of KL signaling because normal expression of *DLK2*, as well as induction by SL or karrikin treatment, depends on MAX2 and KAI2 function (13, 22, 56). We showed that expression of Mp*KAI2B* is positively regulated by the signaling cascade of MpKAI2A and MpMAX2. We also showed that Mp*KAI2B* expression was extensively induced in Mp*smxl* loss-of-function mutants. Expression of *DLK2* is increased in *smxl1,2* double mutants in Arabidopsis (14). This is also the case for MpKAI2B. These similarities between DLK2 and MpKAI2B suggest the possibility that MpKAI2B is the ancestral gene of DLK2. In addition, MAX2-dependent and KAI2-dependent regulation of *DLK2* expression may play a significant role in the fine tuning of this signaling.

In conclusion, we showed that the basic regulatory mechanisms of the proteolysis-dependent signaling pathway containing KAI2, MAX2 and SMXL are conserved in *M. polymorpha*. This implies that this signaling pathway had already arisen in the common ancestors of bryophytes and land plants.

## Materials and Methods

Detailed information about plant materials and growth conditions, plasmid construction, mutagenesis by CRISPR, generation of transgenic *M. polymorpha* plants, RNA extraction and expression analysis, application of *rac-*GR24, KAR_1_ and KAR_2_ and DSF analysis are described in the SI Appendix, SI Materials and Methods.

## Acknowledgments

We thank Ottoline Leyser and Jiayang Li for providing 35Spro:SMXL7-YFP and 35Spro:SMXL6-CFP plasmids, and D53 antibody, respectively. This work was supported by a Grants-in-Aid from the Ministry of Education, Culture, Sports, Science, and Technology, Japan (20H05684, 18K19198, 17H06475 16K14748) to J.K

## Sl Materials and Methods

### Plant Materials and Growth Conditions

Strains Takaragaike-1 (Tak-1; Japanese male line) and Takaragaike-2 (Tak-2; Japanese female line) (1) were used as the WTs in this study. Growth phenotypes were observed using Tak-1. Plants were grown on half-strength Gamborg’s B5 medium with 1.0 % agar under continuous light (50 ~ 60 μmol photons m^−2^s^−1^) at 22 °C.

Accession numbers analyzed in this study were Mp*KAI2A*: Mapoly0023s0137.1, Mp*KAI2B*: Mapoly0031s0148.1, Mp*MAX2*: Mapoly0113s0006.1, Mp*SMXL*: Mapoly0006s0101.1.

### Mutagenesis by CRISPR

pMpGE_En01 (GenBank LC090754; Sugano et al., 2018), digested with *Sac*I and *Pst*I, was separated by electrophoresis and purified using the GE Healthcare illustra™ GFX™ PCR DNA and Gel Band Purification Kit. Primers were treated at 96 °C for 5 min to make them double-stranded (ds). The ds primers were cloned into pMpGE_En01 (2; GenBank LC090756) using the Takara In-Fusion^®^ HD Cloning Kit. The LR reaction was performed using the resultant plasmid (Entry clone) and pMpGE010 (Destination vector (2) to make a plasmid containing gRNA and Cas9.

### Transformation of *M. polymorpha*

Plasmids were transformed into *Agrobacterium tumefaciens* strain GV2260 by electroporation using the GenePulser Xcell™ (BIO-RAD). The transformed *Agrobacterium* was introduced into cut thallus according (3). Selection was performed using hygromycin or chlorsulfuron.

### RNA extraction and expression analysis

RNA was extracted from frozen plant samples using the NucleoSpin RNA Plant kit (MACHEREY-NAGEL). cDNAs were synthesized using SuperScriptIII VILO (Invitrogen). Quantitative PCR was performed using the SYBR Green I with Light Cycler 480 system (Roche Applied Science). Primers used to amplify the cDNAs are shown in Supplemental Table 1. Expression of the endogenous MpSMXL and the introduced Mp*SMXL^d53^* genes were amplified with primer pairs endoMpSMXL-d53_qPCR-F and endoMpSMXL_qPCR-R for Mp*SMXL* and endoMpSMXL-d53_qPCR-F MpSMXL and d53_qPCR-R for the Mp*SMXL^d53^*. The elongation factor gene Mp*EF1* (Mapoly0024s0116.1) and Actin (Mapoly0016s0139.1) were used as standards.

### Analysis of thallus phenotypes

Thallus area was measured using 14-day-old plants. To accurately measure the area, plants were flattened with a cover slip on a glass slide and images were taken. The area of the thallus was analyzed using image processing software (ImageJ). To measure the angle between the thallus and the growth medium, images were taken from the side and analyzed using ImageJ.

### Application of *rac*-GR24, KAR_1_ and KAR_2_

The karrikins KAR_1_ and KAR_2_ and the potent synthetic strigolactone analog *rac-*GR24 were obtained from Chiralix B.V., dissolved in acetone and added to half-strength Gamborg’s B5 medium. For mock treatment, acetone was added to the medium.

### Differential Scanning Fluorimetry (DSF) analysis

The coding sequences for MpKAI2A and MpKAI2B were amplified from cDNA synthesized from total mRNA of the *Marchantia polymorpha* thallus using primers described in Supplemental Information Table 1. The PCR products were cloned into the pHIS8 vector (Addgene) using the In-Fusion HD Cloning Kit (Clontech) to generate pHIS8-MpKAI2A and pET28M-MpKAI2B. BL21 (DE3) (Takara) was used for recombinant protein expression. Overnight cultures (3 mL) in LB liquid medium containing 100 μg/ kanamycin were inoculated into fresh LB medium (1 L) containing 100 μg/ kanamycin and incubated at 37 °C until the OD600 value reached 0.6 ~ 0.8. Then, IPTG was added to 200 μM, and the cells were further incubated at 25 °C overnight. The cells were collected by centrifugation at 4000 rpm for 15 minutes. The pellets were resuspended in 30 mL of buffer A (20 mM Tris-HCl (pH 8.0) and 200 mM NaCl). The supernatants (30 mL) from the resulting lysates were crushed using a cell disruption device. The fractions were purified with Ni affinity beads pre-equilibrated with buffer A. MpKAI2A and MpKAI2B proteins were eluted with 5 mL of buffer A containing 125 mM of imidazole. The purified proteins were concentrated using an Amicon Ultra-4 10 K (Millipore).

For DSF analysis, 20 μl reaction mixtures containing 10 μg protein, 0.015 μl Sypro Orange and GR24s with final concentrations of 250 μM, 100 μM or 50 μM were prepared in 96-well plates. The final acetone concentration in the reaction mixture was 5 %. 1xPBS buffer (pH 7.4) containing 5 % acetone was used in the control reaction. DSF experiments were performed using a Light Cycler 480 II (Roche). Sypro Orange (Invitrogen) was used as the reporter dye. Samples were incubated at 25 °C for 10 min, then, the fluorescence wavelength was detected continuously from 30 °C to 95 °C. The denaturation curve was obtained using MxPro software (Agilent).

---

**Supplementary Information Appendix Fig. S1.**
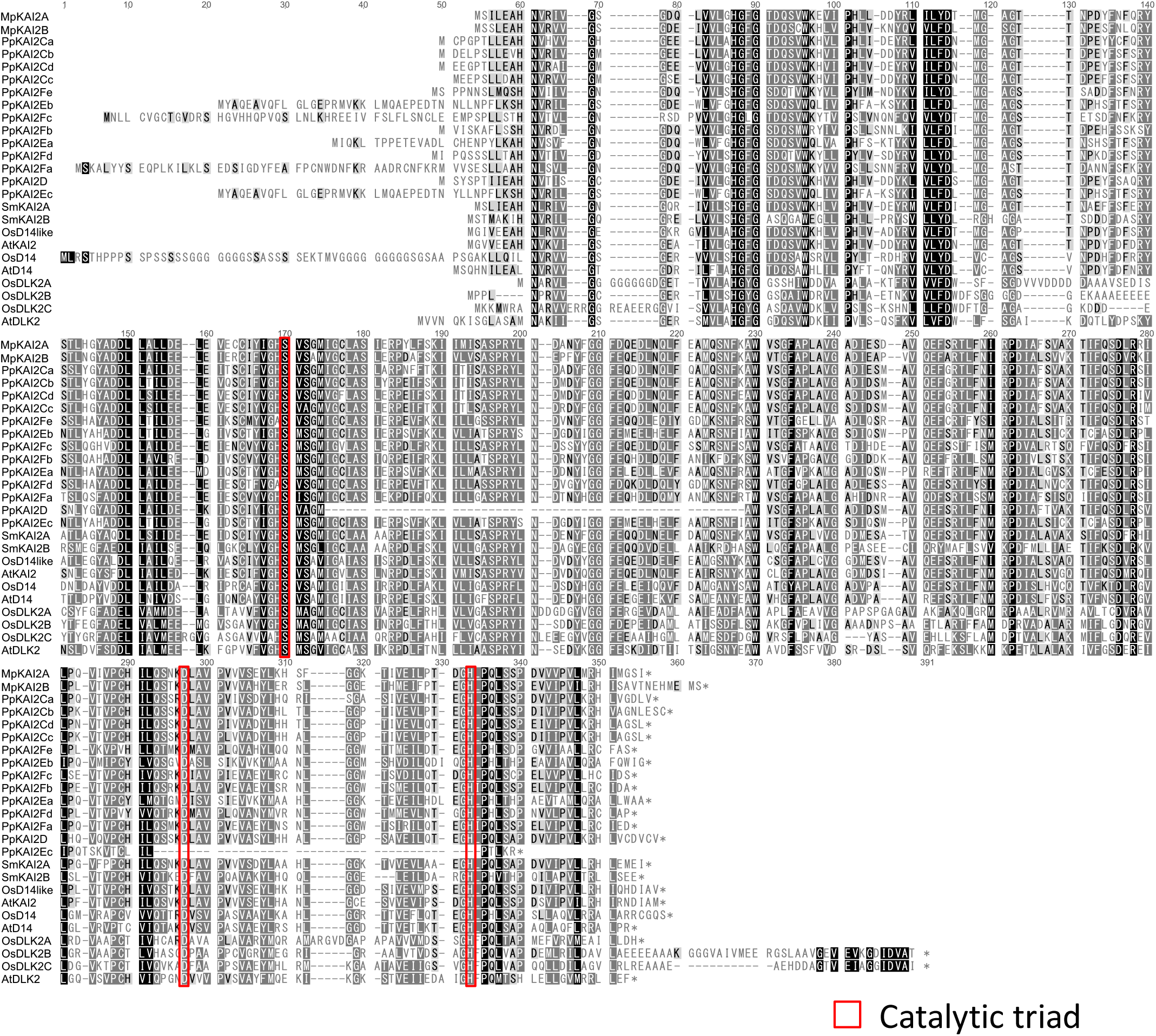
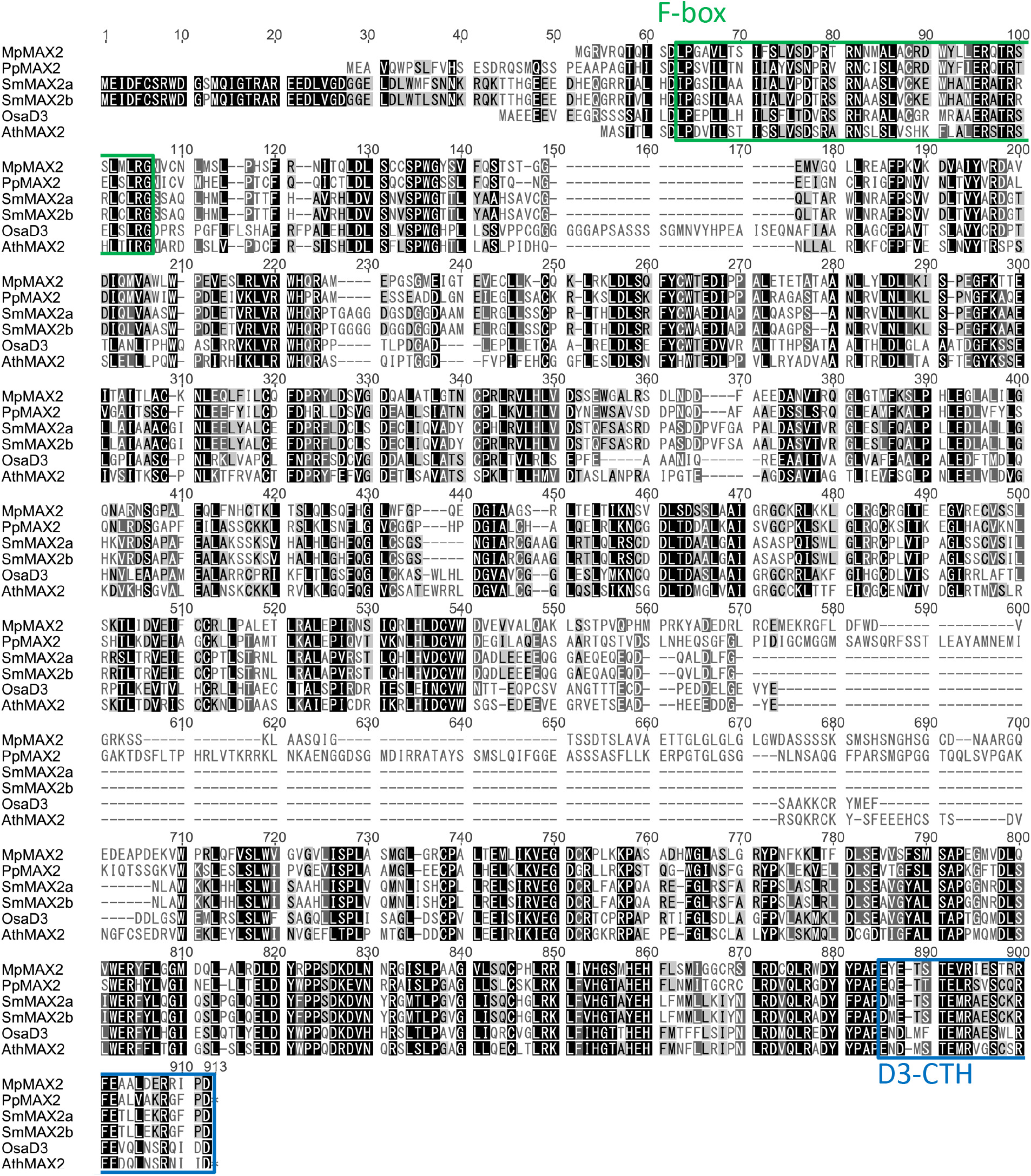
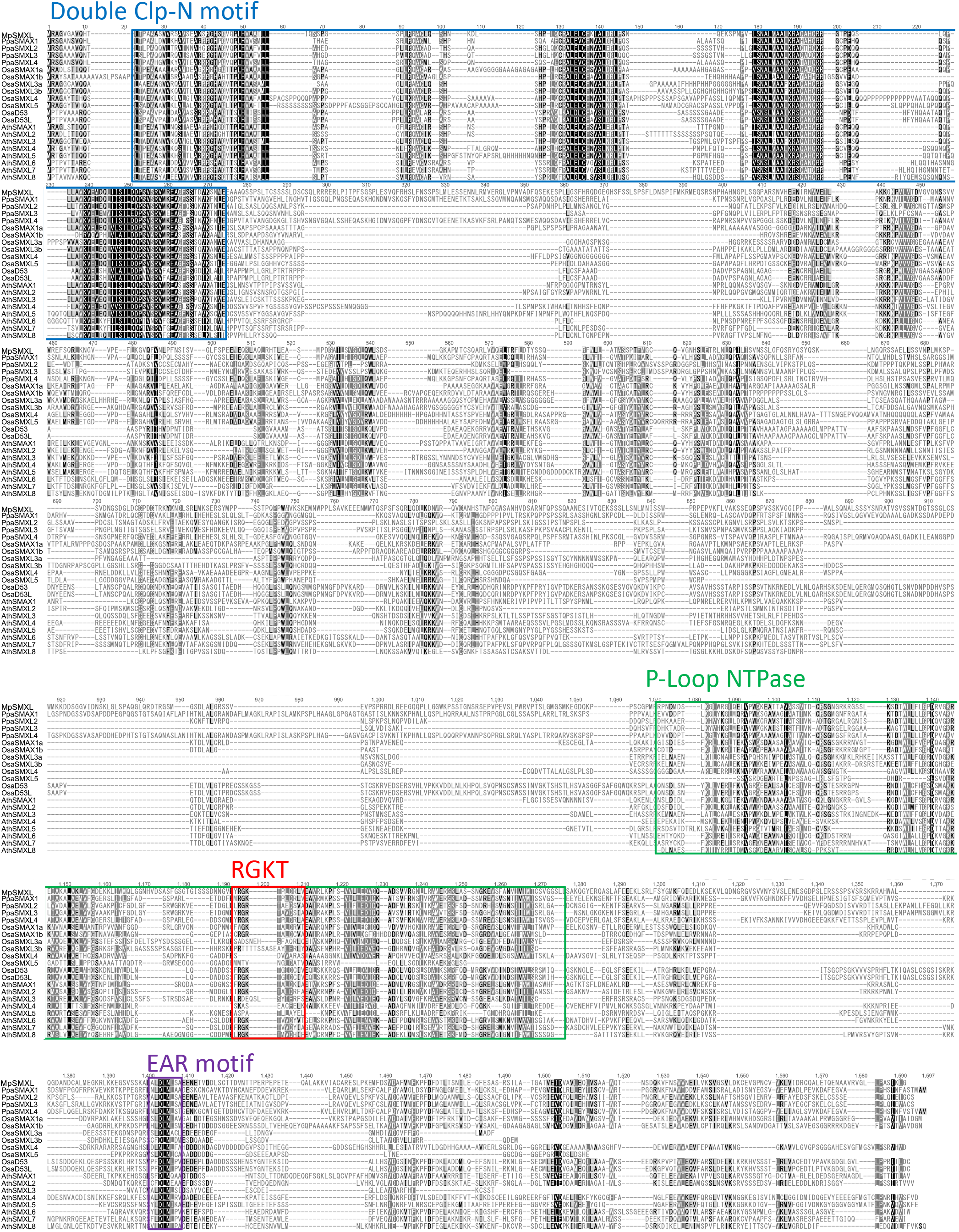
Alignments of amino acid sequences of KAI2 and D14 proteins (A), MAX2 (B) and SMXL proteins (C). Catalytic triad amino acids (S, D and H) conserved in KAI2 and D14 proteins are marked with red squares. Mp, *Marchantia polymorpha*; Pp, *Physcomitrella patens*; Sm, *Selaginella moellendorffii*; Os, *Oryza sativa*; At *Arabidopsis thaliana*; DLK2, DWARF 14-LIKE 2;

**Supplementary Information Appendix Fig. S2.**
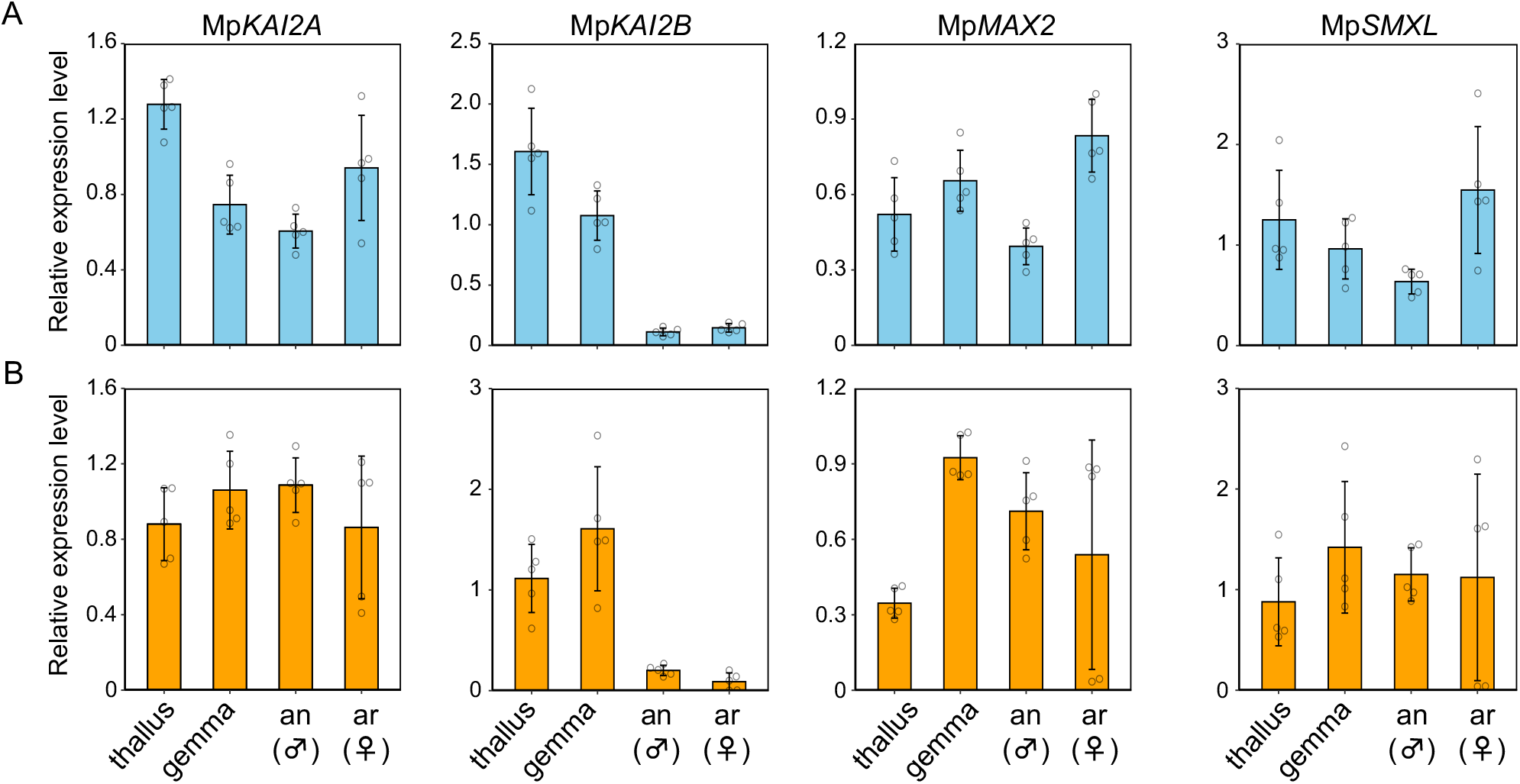
Expression patterns of KL signaling genes. Expression levels of Mp*KAI2A*, Mp*KAI2B*, Mp*MAX2* and Mp*SMXL* relative to the *M. polymorpha ELONGATION FACTOR-1α* gene (Mp3g23400.1) **(A)** and *Actin* gene (Mp6g11010.1) **(B)** in the thallus, gemma, antheridiophore (an) and archegoniophore (ar) are shown. qPCR was used to measure expression levels. Data are the mean ± SD from five biological replications.

**Supplementary Information Appendix Fig. S3.**
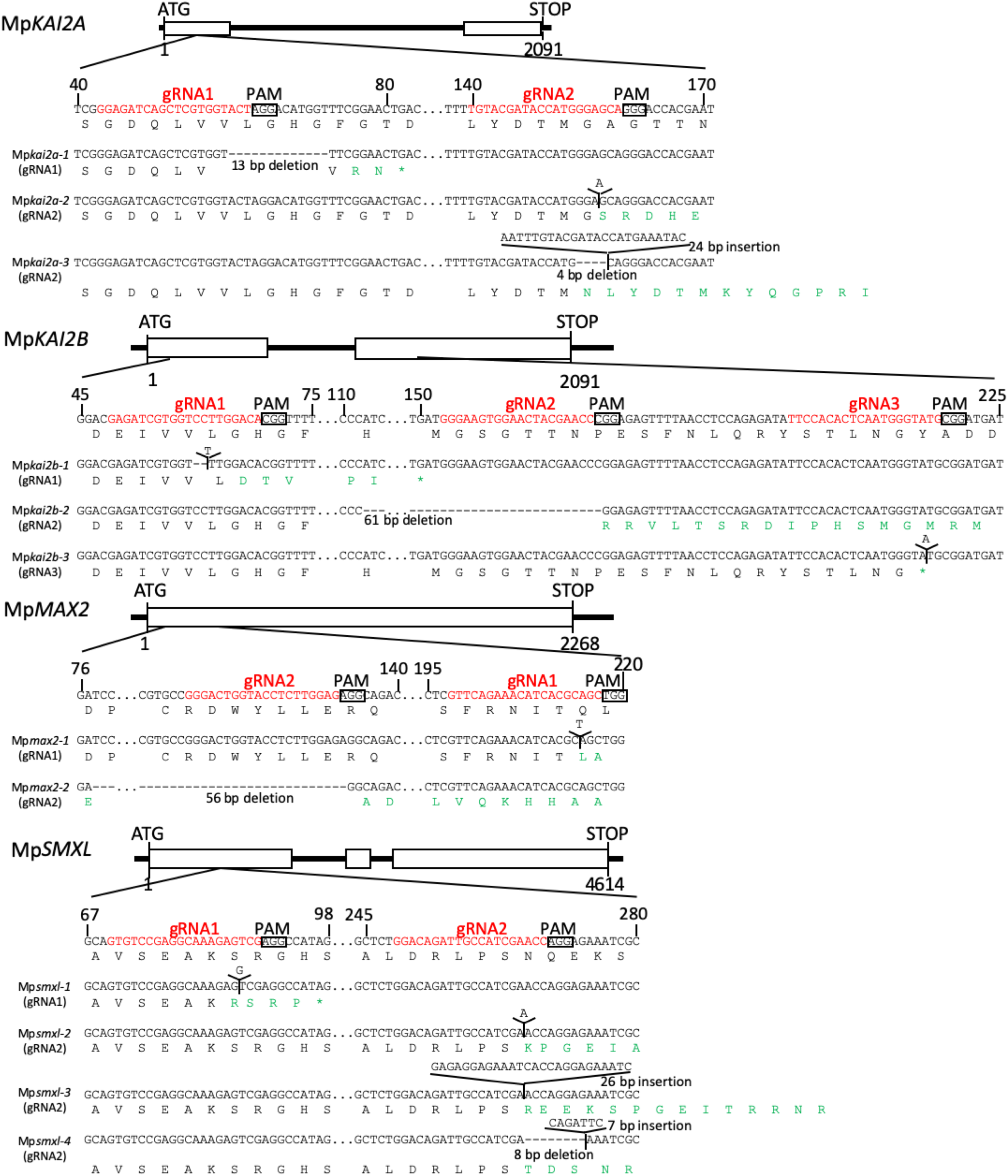
Production of CRISPR mutants of KL signaling genes. Structure of genes, position of guide RNAs (gRNA)s, PAM sequences and resultant mutations are shown. Predicted amino acid sequences in WT proteins are shown in black and those in mutants are shown in green letters. Asterisks indicate stop codons.

**Supplementary Information Appendix Fig. S4.**
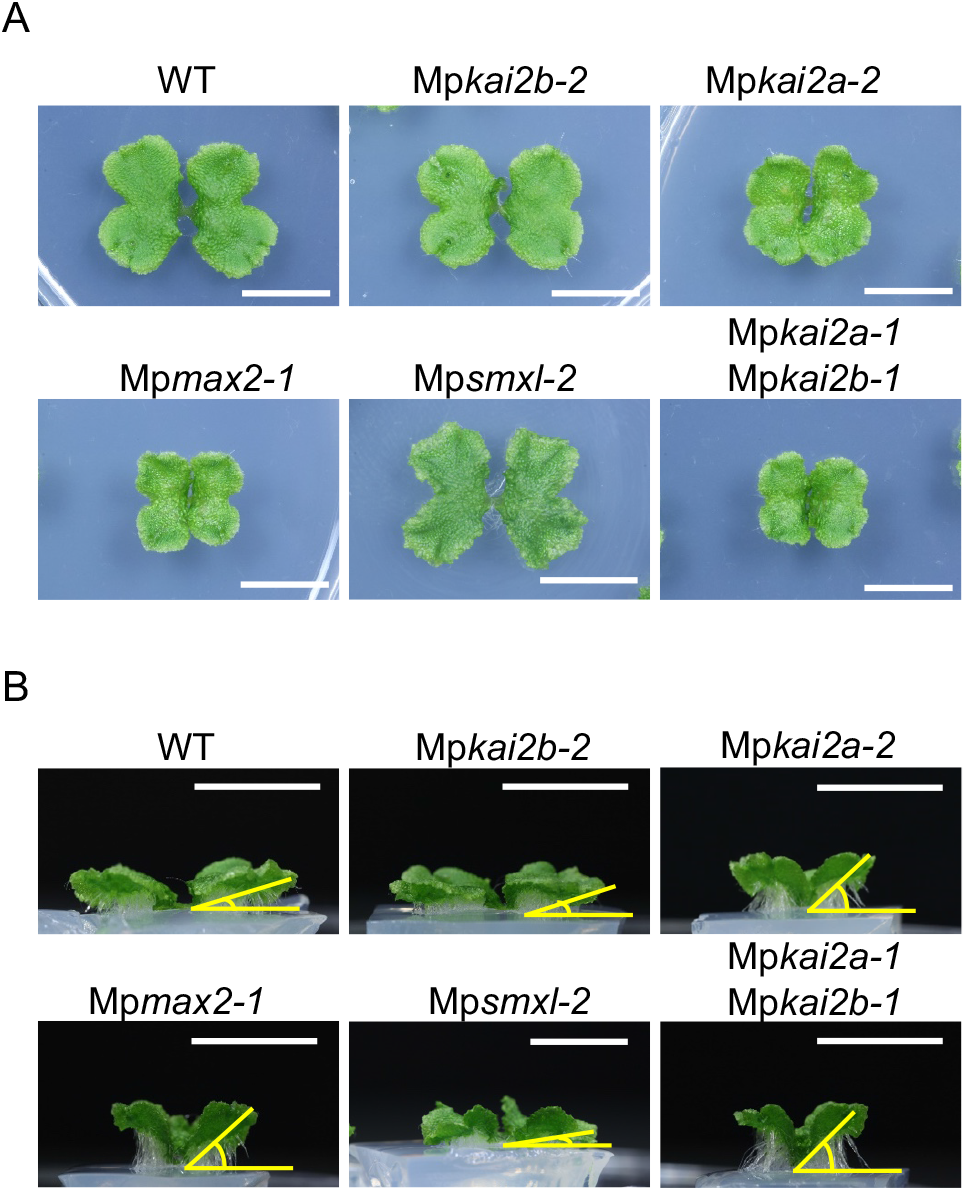
Growth phenotypes of loss-of-function mutants of KL signaling genes. Top **(A)** and side **(B)** views of 14-day-old thalli of Mp*kai2a-2*, Mp*kai2b-2*, Mp*max2-1* and Mp*smxl-2*, and Mp*kai2a-1* Mp*kai2b-1*mutants. Bars = 1 cm.

**Supplementary Information Appendix Fig. S5.**
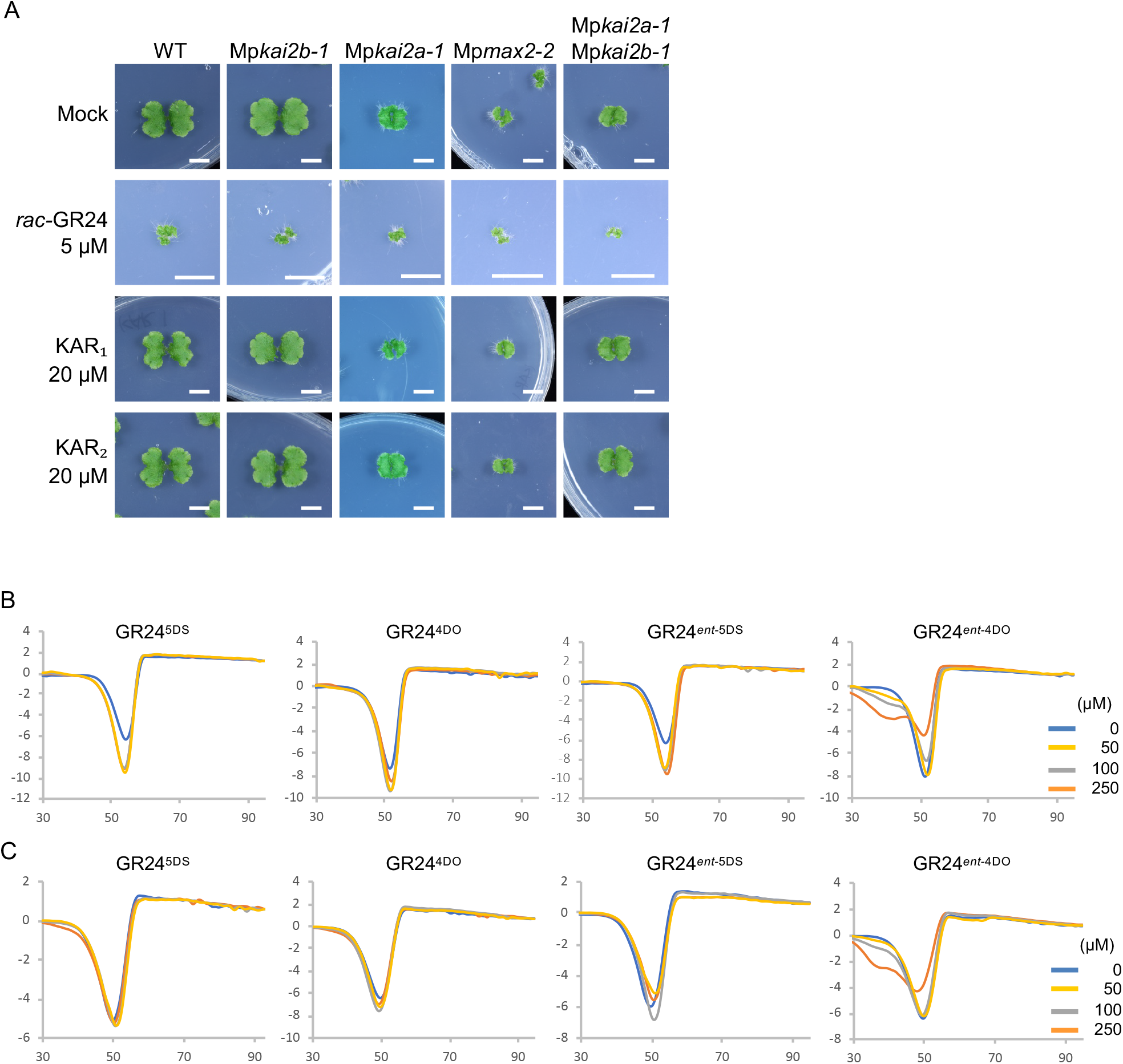
Effects of exogenously supplied *rac*-GR24, KAR_1_, KAR_2_ on thallus growth. Top view **(A)** of thalli grown on the media containing *rac*-GR24, KAR_1_ or KAR_2_. Bars = 1 cm. Differential Scanning Fluorimetry (DSF) analysis with 4 stereoisomers of GR24 and MpKAI2A **(B)** and MpKAI2B **(C)** proteins. A slight change of melting points was detected when the GR24^*ent*-5DS^ was incubated with MpKAI2A and MpKAI2B. The enantiomers *ent*-5DS and *ent*-4DO have not been reported to be produced in nature.

**Supplemental Table 1:**
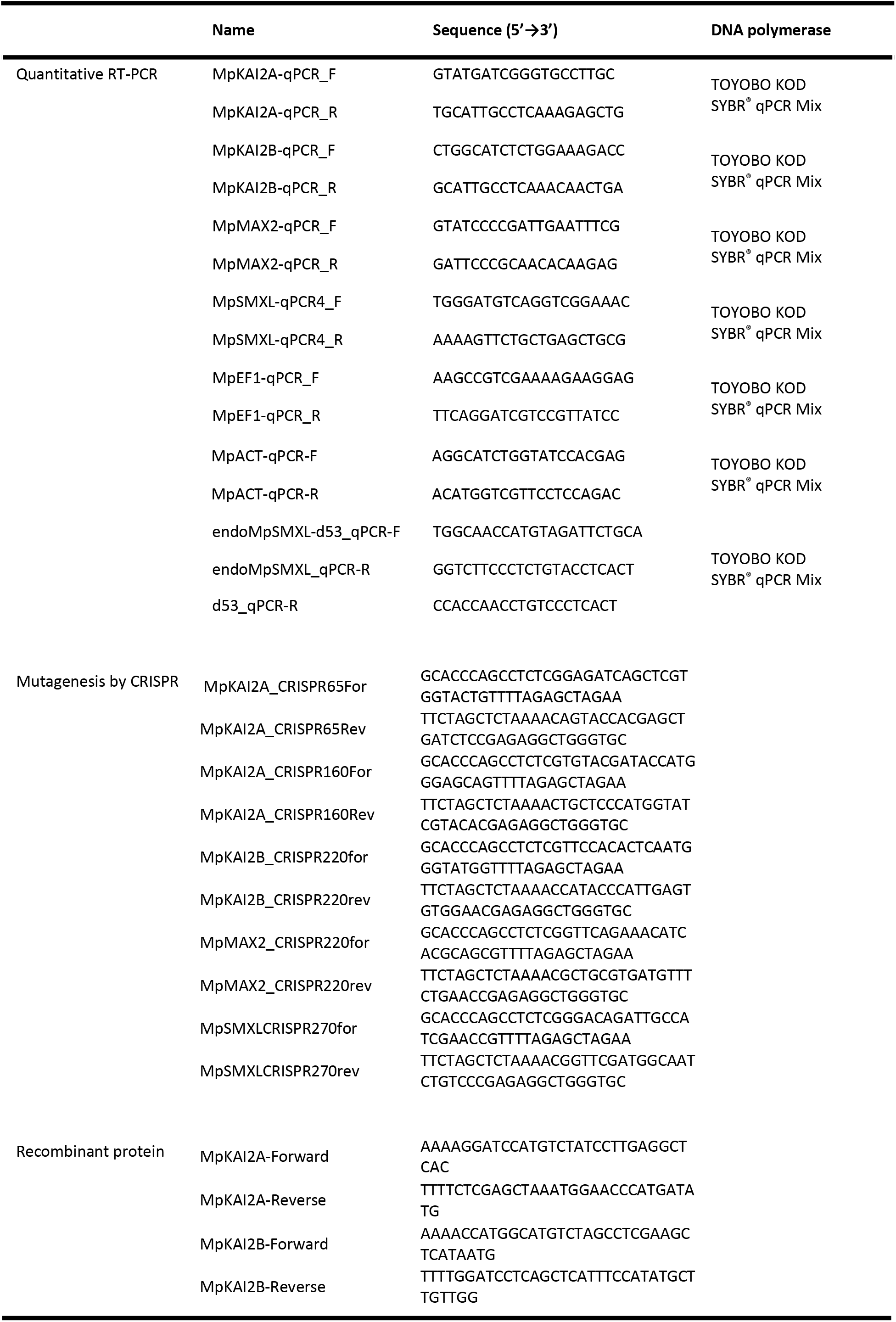
Primers used in this study

**Supplemental Table 2:** Gene Accession

|  | Name | Accession |  | Name | Accession |
| --- | --- | --- | --- | --- | --- |
| KAI2A/B | MpKAI2A | Mp2g11710.1 | SMXL | MpSMXL | Mp3g06310.1 |
|  | MpKAI2B | Mp2g04930.1 |  | PpSMAX1 | XP_024395920.1 |
|  | PpKAI2Ca | XP_024394426.1 |  | PpSMXL2 | XP_024386397.1 |
|  | PpKAI2Cb | XP_024365703.1 |  | PpSMXL3 | XP_024368357.1 |
|  | PpKAI2Cc | XP_024376426.1 |  | PpSMXL4 | XP_024384104.1 |
|  | PpKAI2Cd | XP_024379297.1 |  | OsaSMAX1a | XP_015650285.1 |
|  | PpKAI2D | XP_024379297.1 |  | OsaSMAX1b | XP_015623668.1c |
|  | PpKAI2Ea | XP_024374275.1 |  | OsaSMXL3a | XP_015615327.1 |
|  | PpKAI2Eb | XP_024372547.1 |  | OsaSMXL3b | XP_015636646.1 |
|  | PpKAI2Ec | XP_024366375.1 |  | OsaSMXL4 | XP_015636167.1 |
|  | PpKAI2Fa | XP_024373000.1 |  | OsaSMXL5 | XP_025878695.1 |
|  | PpKAI2Fb | XP_024390568.1 |  | OsaD53 | XP_015617462.1 |
|  | PpKAI2Fc | XP_024369665.1 |  | OsaD53L | XP_015620371.1 |
|  | PpKAI2Fd | XP_024368082.1 |  | AthSMAX1 | NP_200579.1 |
|  | PpKAI2Fe | XP_024386103.1 |  | AthSMXL2 | NP_194764.1 |
|  | SmKAI2A | XP_024533478.1 |  | AthSMXL3 | NP_190817.1 |
|  | SmKAI2B | XP_002969138.1 |  | AthSMXL4 | NP_194721.1 |
|  | OsD14like | XP_015617518.1 |  | AthSMXL5 | NP_568849.2 |
|  | AtKAI2 | NP_195463.1 |  | AthSMXL6 | NP_001077474.1 |
|  | OsD14 | XP_015631400.1 |  | AthSMXL7 | NP_565689.1 |
|  | AtD14 | NP_566220.1 |  | AthSMXL8 | NP_973646.1 |
|  | OsDLK2A | XP_015638523.1 |  |  |  |
|  | OsDLK2B | XP_015650138.1 |  |  |  |
|  | OsDLK2C | XP_015638017.1 |  |  |  |
|  | AtDLK2 | NP_189085.1 |  |  |  |
| MAX2 | MpMAX2 | Mp1g02580.1 |  |  |  |
|  | PpMAX2 | Pp3c17_1180V3.1 |  |  |  |
|  | SmMAX2a | XP_002968649.2 |  |  |  |
|  | SmMAX2b | XP_002989898.2 |  |  |  |
|  | OsaD3 | BAD69288.1 |  |  |  |
|  | AthMAX2 | NP_565979.1 |  |  |  |

